# Partial genome characterization of a novel potentially zoonotic parapoxvirus in a horse, Finland

**DOI:** 10.1101/2023.03.21.533517

**Authors:** Jenni Virtanen, Maria Hautaniemi, Lara Dutra, Ilya Plyusnin, Katja Hautala, Teemu Smura, Olli Vapalahti, Tarja Sironen, Ravi Kant, Paula M. Kinnunen

## Abstract

We report a sequencing protocol and 121 kb poxvirus sequence from a clinical sample of a horse with dermatitis. Based on phylogenetic analyses, the virus is a novel parapoxvirus. We show association with recent epidemic, and previous data suggest zoonotic potential. Further characterization and development of diagnostic protocols are necessary.

## Text

Parapoxviruses (PPVs) usually cause contagious skin infections in ruminants and occasionally infect other species such as humans (*1*). The genus *Parapoxvirus* currently encompasses the following recognized species: Orf virus (ORFV), Bovine Papular Stomatitis Virus (BPSV), Pseudocowpoxvirus (PCPV), Red deerpox virus (RDPV), and Grey sealpox virus (GSEPV) (*2*). Of these, ORFV, BPSV, PCPV, and RDPV are famously zoonotic. PPV genomes are usually 130-140 kb (*2*). Recently, poxviruses have emerged in humans and horses *(3, 4)*.

A severe infection caused by a parapox-like virus (F14.1158H) was first verified from a horse euthanized in Finland in 2013 (*5*). According to the short sequences (1.1 kb in total) obtained from envelope phospholipase (ORF011) and RNA polymerase subunit RPO147 (ORF056) genes, F14.1158H is most closely related to PPVs and is similar to the 585 bp sequences detected in lesions from humans after contact with horses and donkeys in the United States *(5, 6)*. However, the actual classification remained unclear due to limited sequence data and lack of amplification in numerous PPV-PCR assays (*5*).

No other clinical cases were confirmed until 2022 when an epidemic of dermatitis emerged in horses across Finland. Parapoxvirus infection was subsequently identified in several cases using pan-PPV-PCR (*7*) and Sanger sequencing (Appendix). Partial ORF011 sequences were 97% identical to the 2013 case with identity of 79-87% to other parapoxviruses (Table A1). This highlighted the need to properly characterise F14.1158H.

To better characterise the virus, DNA extracted directly from a skin lesion of the 2013 equine case (*5*) was subjected to next-generation sequencing (NGS) with two different protocols (Appendix). First protocol relying on a pool of poxvirus primers was insufficient to acquire enough sequence data. With a PCR-free approach, utilising enrichment of the viral DNA, we acquired as much as 121 kb of nucleotide sequence, almost full genome, with coverage values of approximately 100 in 5 contigs (BioProject: PRJNA922554 and GenBank: OQ248663-OQ248667). Overall GC-content was 68.4%, which is similar to PPVs (*2*). Due to the lack of a reference genome, the data could not be fully assembled and oriented. The lack of high-quality DNA and unsuccessful virus isolation attempts (Appendix) further complicated the sequencing process. In the future, it is important to fill the gaps to ensure finalization of a complete genome that can be used as a reference. In another study, a combination of short- and long-read sequencing was used to recover the full genome of the GSEPV (*8*). However, with our clinical sample, the small amount of DNA available for sequencing led to an alternative approach.

Phylogenetic analysis was conducted for the following poxvirus core genes (*9*) (ORF numbers designated according to PPV ORFs (*10*)): DNA polymerase (ORF025), early transcription factor VETFL (ORF083), DNA-directed RNA polymerase subunit RPO132 (ORF101), and DNA topoisomerase type 1 (ORF062) (Figure). Consistent with earlier observations based on partial sequences of ORF11 and ORF056, F14.1158H grouped clearly closer to PPVs than other poxviruses (Figure A), although distinctly separate from the five currently recognized species (Figure B-D). Amino acid sequences were more similar to PPV species than to other Chordopoxviruses. For example, amino acid identity of DNA polymerase was 46-60% between F14.1158H and viruses from other genera, 76-80% between F14.1158H and PPVs, and 84-95% between the previously recognized PPV species (Table A3). Within the PPVs, F14.1158H generally showed the second lowest pairwise nucleotide identity of the group (after the most divergent GSEPV) (Table) with identities to other PPVs being 74-83% (ORF025), 73-83% (ORF083), 78-87% (ORF101), and 84-91% (ORF062). GSEPV was consistently furthest from F14.1158H whereas F14.1158Hs identities to other PPVs were similar. A relatively high difference explains why F14.1158H was not detected by several PCRs designed for detecting PPV, which should be considered when designing diagnostic protocols. These phylogenetic results and sequence identities together with the high GC-content and disease characteristics indicate that F14.1158H is a novel PPV, Equine parapoxvirus (EqPPV). The final taxonomic position and the possible differences of human and equine-derived variants (*6*) will require more data.

**Figure. A).**
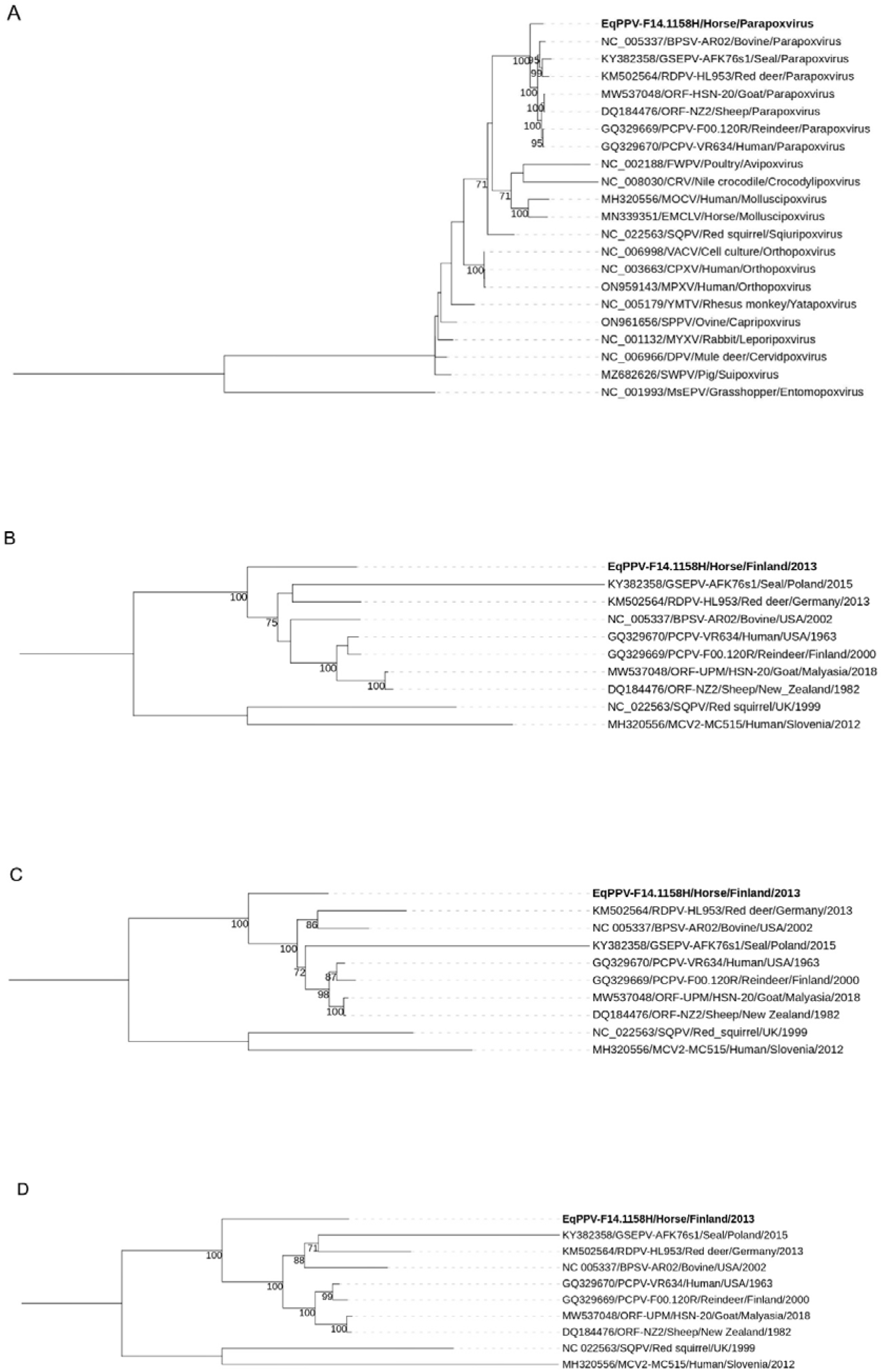
Grouping of F14.1158H among all the genera of the subfamily *Chordopoxvirinae* in a phylogenetic tree based on amino acid sequences of the DNA polymerase (ORF025) gene. B-D) Grouping of F141158H among the genus Parapoxvirus in phylogenetic trees based on the nucleotide sequences of the early transcription factor (ORF083) (B), RNA polymerase (ORF101) (C), and topoisomerase 1 (ORF062) (D) genes. CRV = Crocodilepox virus, EMCLV = Equine molluscum contagiosum-like virus, MOCV = Molluscum contagiosum virus, SQPV = Squirrelpox virus, CPXV = Cowpox virus, MPXV = Monkeypox virus, VACV = Vaccinia virus, YMTV = Yaba monkey tumor virus, SPPV = Sheeppox virus, MYXV = Myxoma virus, SWPV = Swinepox virus, DPV = Deerpox virus, FWPV = Fowlpox virus, MsEPV = Melanoplus sanguinipes entomopoxvirus. MsEPV is used as an outgroup in A and SQPV and MOCV in B-D. Bootstrap values above 70% are shown next to the nodes.

Taken together, most known PPVs are zoonotic, and any novel virus detected in animals should be treated with concern (*6*). Thus, considering the tendency of PPVs to cause disease, such as farmyard pox, in humans, EqPPV has a zoonotic potential. Hence, it is important to sample humans and other animals in contact with infected horses. It is also important to establish diagnostic protocols due to low specificity and sensitivity of pan-PPV-PCR for EqPPV (Appendix). Evidently, this virus is of veterinary importance posing a threat for horses and causing financial losses to owners. The information provided here will inform development of proper diagnostic tools enabling also establishment prevention measures.

## Acknowledgements

We thank DVM, PhD, Dipl. ECVP Niina Airas for her expert support regarding equine medicine. We also thank Mira Utriainen for technical assistance.

This study was financially supported by the Niemi Foundation, Finnish Foundation of Veterinary Research, and the Erkki Rajakoski Fund of Hippos Finland.

## About the author

Ms. Virtanen is a post-doctoral researcher at the University of Helsinki in the field of clinical microbiology. Her research interests include zoonotic viruses and pathogens in human-animal interface.

## Conflict of interest declaration

Besides holding the title of Adjunct Professor (Docent) of University of Helsinki, PMK is an employee of MSD Animal Health. This study was initiated before her joining the company. Other authors have nothing to declare.

---

**Table:**
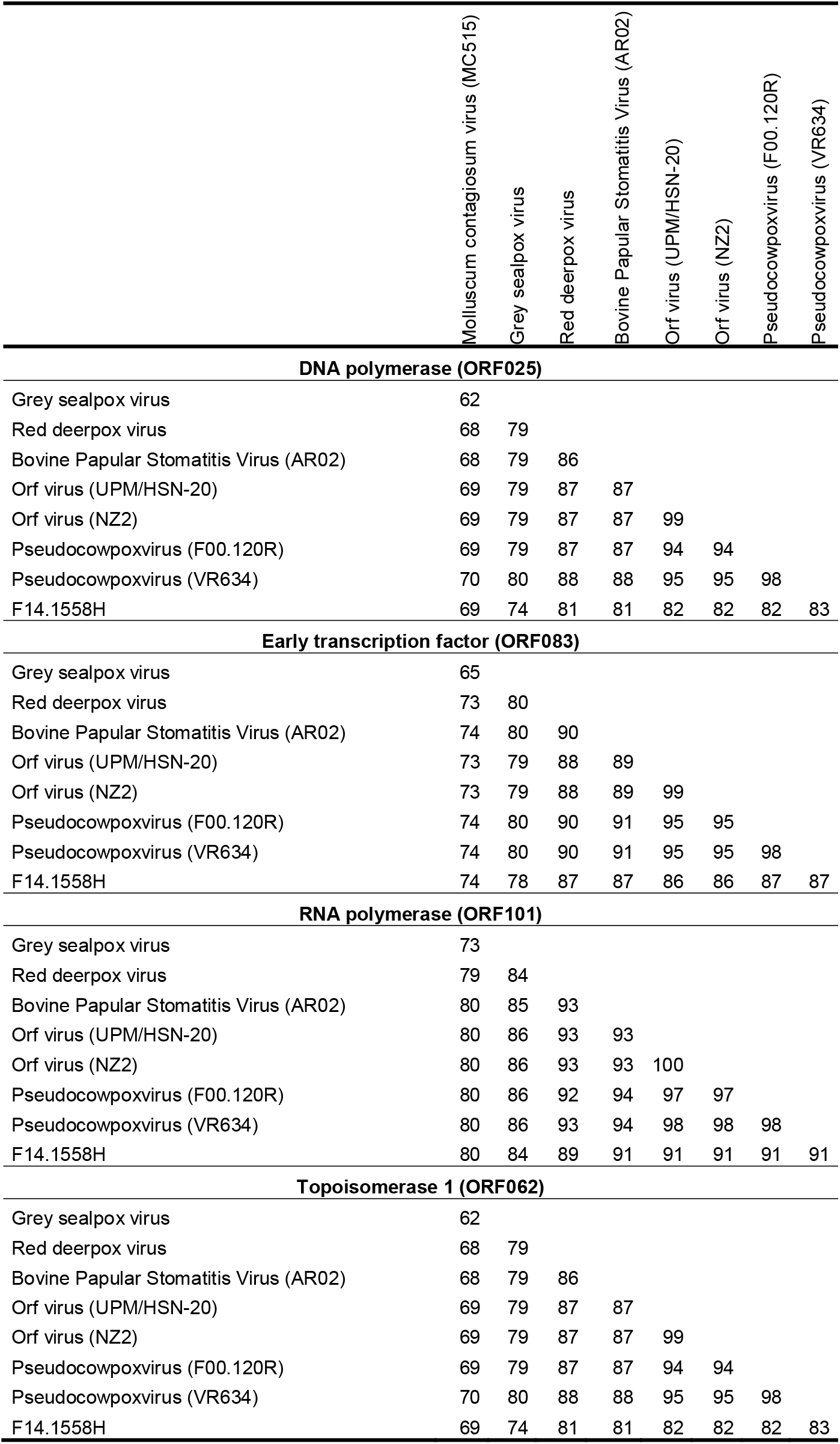
Nucleotide identity (as percentages) comparison between F14.1158H and other parapoxviruses and Molluscum contagiosum virus in the four selected core genes.

|  | Molluscum contagiosum virus (MC515) | Grey sealpox virus | Red deerpox virus | Bovine Papular Stomatitis Virus (AR02) | Orf virus (UPM/HSN-20) | Orf virus (NZ2) | Pseudocowpoxvirus (F00.120R) | Pseudocowpoxvirus (VR634) |
| --- | --- | --- | --- | --- | --- | --- | --- | --- |
| **DNA polymerase (ORF025)** |  |  |  |  |  |  |  |  |
| Grey sealpox virus | 62 |  |  |  |  |  |  |  |
| Red deerpox virus | 68 | 79 |  |  |  |  |  |  |
| Bovine Papular Stomatitis Virus (AR02) | 68 | 79 | 86 |  |  |  |  |  |
| Orf virus (UPM/HSN-20) | 69 | 79 | 87 | 87 |  |  |  |  |
| Orf virus (NZ2) | 69 | 79 | 87 | 87 | 99 |  |  |  |
| Pseudocowpoxvirus (F00.120R) | 69 | 79 | 87 | 87 | 94 | 94 |  |  |
| Pseudocowpoxvirus (VR634) | 70 | 80 | 88 | 88 | 95 | 95 | 98 |  |
| F14.1558H | 69 | 74 | 81 | 81 | 82 | 82 | 82 | 83 |
| **Early transcription factor (ORF083)** |  |  |  |  |  |  |  |  |
| Grey sealpox virus | 65 |  |  |  |  |  |  |  |
| Red deerpox virus | 73 | 80 |  |  |  |  |  |  |
| Bovine Papular Stomatitis Virus (AR02) | 74 | 80 | 90 |  |  |  |  |  |
| Orf virus (UPM/HSN-20) | 73 | 79 | 88 | 89 |  |  |  |  |
| Orf virus (NZ2) | 73 | 79 | 88 | 89 | 99 |  |  |  |
| Pseudocowpoxvirus (F00.120R) | 74 | 80 | 90 | 91 | 95 | 95 |  |  |
| Pseudocowpoxvirus (VR634) | 74 | 80 | 90 | 91 | 95 | 95 | 98 |  |
| F14.1558H | 74 | 78 | 87 | 87 | 86 | 86 | 87 | 87 |
| **RNA polymerase (ORF101)** |  |  |  |  |  |  |  |  |
| Grey sealpox virus | 73 |  |  |  |  |  |  |  |
| Red deerpox virus | 79 | 84 |  |  |  |  |  |  |
| Bovine Papular Stomatitis Virus (AR02) | 80 | 85 | 93 |  |  |  |  |  |
| Orf virus (UPM/HSN-20) | 80 | 86 | 93 | 93 |  |  |  |  |
| Orf virus (NZ2) | 80 | 86 | 93 | 93 | 100 |  |  |  |
| Pseudocowpoxvirus (F00.120R) | 80 | 86 | 92 | 94 | 97 | 97 |  |  |
| Pseudocowpoxvirus (VR634) | 80 | 86 | 93 | 94 | 98 | 98 | 98 |  |
| F14.1558H | 80 | 84 | 89 | 91 | 91 | 91 | 91 | 91 |
| **Topoisomerase 1 (ORF062)** |  |  |  |  |  |  |  |  |
| Grey sealpox virus | 62 |  |  |  |  |  |  |  |
| Red deerpox virus | 68 | 79 |  |  |  |  |  |  |
| Bovine Papular Stomatitis Virus (AR02) | 68 | 79 | 86 |  |  |  |  |  |
| Orf virus (UPM/HSN-20) | 69 | 79 | 87 | 87 |  |  |  |  |
| Orf virus (NZ2) | 69 | 79 | 87 | 87 | 99 |  |  |  |
| Pseudocowpoxvirus (F00.120R) | 69 | 79 | 87 | 87 | 94 | 94 |  |  |
| Pseudocowpoxvirus (VR634) | 70 | 80 | 88 | 88 | 95 | 95 | 98 |  |
| F14.1558H | 69 | 74 | 81 | 81 | 82 | 82 | 82 | 83 |

## Notes

### Summary of Updates

Data has been added concerning the 2022 equine pastern dermatitis outbreak in Finland. Due to this, one author was added. Manuscript language has also been revised.

