## Appendix for "Partial genome characterization of a novel potentially zoonotic parapoxvirus in a horse, Finland"

Testing of the samples from the 2021-2022 outbreak

Skin lesion swab samples from 25 horses suffering from vesicular or other kind of acute dermatitis in the pastern area were collected by veterinarians at ten stables in Finland between December 2021 and March 2022. Given the history of circulation of PPV in horses in Finland *(1),* the samples were tested for the presence of the equine PPV. Swabs were incubated in 500 µl of Dulbecco’s phosphate buffered saline + 0.2% bovine serum albumin in +4°C o/n and DNA was extracted using Viral RNA mini kit (Qiagen) according to manufacturer’s instructions. Samples were then tested with pan-PPV-PCR *(2).* PCR products were run in 1.5% agarose gel. Products of correct size were purified with PCR purification kit or Gel extraction kit (GeneJET) and sequenced with Sanger sequencing.

PCR products from a total of 16 horses were sequenced, and poxvirus DNA was confirmed from nine (56%) of these. The rest were either host genome or had too many overlapping sequences to enable the confirmation of the presence or absence of poxvirus sequences. Sanger sequences from five horses from three stables had good enough quality for further analyses. These five sequences were compared with the ones obtained from the 2013 case (PPV variant F14.1158H; *(1))* and selected strains from GenBank with phylogenetic analyses and nucleotide identity calculations (Figure A1 and Table A1, methods described in ‘PCR-free sequencing and sequence analysis’ -section). Sequences of 2022 outbreak were 99-100% identical to each other based on partial envelope phospholipase gene (ORF011) and 97% identical to the sequence of the 2013 case. They were only 79-87% identical to other PPVs and were hence concluded to most likely represent the same PPV species, which infected the index case in 2013. Sequences were deposited in GenBank with accession numbers OR112284-OR112288.

The pan-PPV-PCR showed poor specificity to equine PPV in horse skin samples. Furthermore, due to the sequence difference of EqPPV with the other PPVs (Table A1), we cannot trust the sensitivity either. Thus, the remaining 16 samples, from which we did not obtain clear PCR results or sequences, can’t be considered negative. The obvious unspecificity results at least partially from host DNA in PCR products. It is important to develop more reliable diagnostic protocols to prepare for further epidemics.

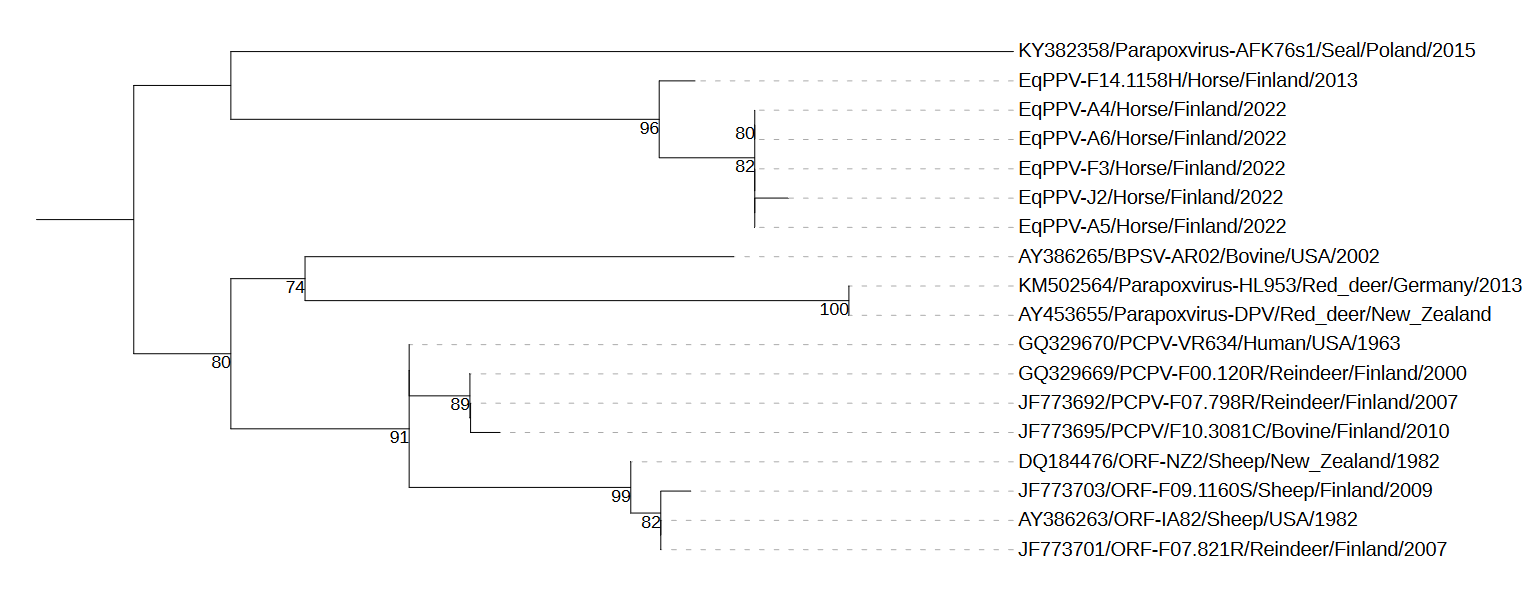

Figure A1: Phylogenetic tree of partial envelope phospholipase gene (ORF011, bp 449-620) including F14.1158, five sequences from 2022 pastern dermatitis outbreak, and selected representatives of other parapoxvirus species. Bootstrap values above 70% are shown next to the nodes.

Table A1: Nucleotide identities as percentages between sequences from Figure S1. Identities within species are highlighted with gray.

|  | EqPPV-A5/Horse/Finland/2022 | EqPPV-A6/Horse/Finland/2022 | EqPPV-A4/Horse/Finland/2022 | EqPPV-J2/Horse/Finland/2022 | EqPPV-F3/Horse/Finland/2022 | EqPPV-F14.1158H/Horse/Finland/2013 | JF773701/ORF-F07.821R/Reindeer/Finland/2007 | AY386263/ORF-IA82/Sheep/USA/1982 | GQ329669/PCPV-F00.120R/Reindeer/Finland/2000 | DQ184476/ORF_NZ2/Sheep/New_Zealand/1982 | AY386265/BPSV-AR02/Bovine/USA/2002 | JF773695/PCPV/F10.3081C/Bovine/Finland/2010 | GQ329670/PCPV-VR634/Human/USA/1963 | JF773703/ORF-F09.1160S/Sheep/Finland/2009 | JF773692/PCPV-F07.798R/Reindeer/Finland/2007 | AY453655/Parapoxvirus-DPV/Red_deer/New_Zealand | KM502564/Parapoxvirus-HL953/Red_deer/Germany/2013 |
| --- | --- | --- | --- | --- | --- | --- | --- | --- | --- | --- | --- | --- | --- | --- | --- | --- | --- |
| EqPPV-A6 (2022) | 100 |  |  |  |  |  |  |  |  |  |  |  |  |  |  |  |  |
| EqPPV-A4 (2022) | 100 | 100 |  |  |  |  |  |  |  |  |  |  |  |  |  |  |  |
| EqPPV-J2 (2022) | 99 | 99 | 99 |  |  |  |  |  |  |  |  |  |  |  |  |  |  |
| EqPPV-F3 (2022) | 100 | 100 | 100 | 99 |  |  |  |  |  |  |  |  |  |  |  |  |  |
| EqPPV-F14.1158H (2013) | 97 | 97 | 97 | 97 | 97 |  |  |  |  |  |  |  |  |  |  |  |  |
| JF773701/ORF-F07.821R | 84 | 85 | 85 | 84 | 85 | 84 |  |  |  |  |  |  |  |  |  |  |  |
| AY386263/ORF-IA82 | 84 | 86 | 86 | 84 | 85 | 84 |  |  |  |  |  |  |  |  |  |  |  |
| GQ329669/PCPV-F00.120R | 83 | 85 | 85 | 83 | 84 | 85 | 94 | 94 |  |  |  |  |  |  |  |  |  |
| DQ184476/ORF_NZ2 | 85 | 87 | 87 | 85 | 86 | 84 | 99 | 99 | 95 |  |  |  |  |  |  |  |  |
| AY386265/BPSV-AR02 | 82 | 82 | 82 | 82 | 82 | 83 | 88 | 86 | 88 | 87 |  |  |  |  |  |  |  |
| JF773695/PCPV/F10.3081C | 83 | 85 | 85 | 83 | 84 | 85 | 94 | 94 | 99 | 94 | 87 |  |  |  |  |  |  |
| GQ329670/PCPV-VR634 | 84 | 86 | 86 | 84 | 85 | 86 | 95 | 95 | 99 | 96 | 89 | 98 |  |  |  |  |  |
| JF773703/ORF-F09.1160S | 84 | 85 | 85 | 84 | 84 | 83 | 99 | 99 | 94 | 99 | 86 | 93 | 95 |  |  |  |  |
| JF773692/PCPV-F07.798R | 83 | 85 | 85 | 83 | 84 | 85 | 94 | 94 | 100 | 95 | 88 | 100 | 99 | 94 |  |  |  |
| AY453655/Parapoxvirus-DPV | 80 | 81 | 81 | 79 | 80 | 82 | 83 | 84 | 87 | 85 | 85 | 87 | 87 | 84 | 87 |  |  |
| KM502564/Parapoxvirus-HL953 | 80 | 82 | 82 | 79 | 81 | 81 | 83 | 85 | 87 | 85 | 86 | 87 | 88 | 85 | 87 | 100 |  |
| KY382358/Parapoxvirus-AFK76s1 | 82 | 83 | 83 | 81 | 82 | 82 | 80 | 80 | 82 | 81 | 80 | 82 | 84 | 80 | 83 | 80 | 80 |

Sequencing using a primer pool

DNA (a total of 4 µg at a concentration of 215 ng/µl) had been extracted from a skin lesion of an infected horse in our earlier study (*1*). In the first attempt to obtain enough sequence data for proper characterization from a very limited amount of sample, we used an enrichment PCR with a combination of poxvirus primers and next generation sequencing (NGS). PCR reactions contained 23.5 µl of water, 2 µl of the sample, 10 µl Taq buffer (NH_4_)_2_SO_4_, 1 µl of 10 mM dNTP, 4 µl of 25 mM MgCl_2_, 0.5 µl of Taq DNA polymerase (Thermo Scientific), and 1 µl of primer pool including 10 µM of each primer. The program was as follows: 95˚C for 1 min, and 30 rounds of denaturation (95˚C 30 s), annealing (50˚C 30 s) and extension (72˚C 4 min), followed by final extension (72˚C 10 min). Two PCR reactions with different primer combinations were run: one using the combination of primers listed in Table A2 and another with reverse complementary sequences of the same primers. The aim was to amplify both the sequences flanked by the primers and the sequences outside the designed product by using the reverse complement. PCR products were combined and purified with GeneJET PCR purification kit (Thermo Scientific).

Libraries were prepared with Nextera XT DNA Sample preparation and Nextera XT Index kits (Illumina). Sequencing was performed with MiSeq (Illumina) and MiSeq reagent kit V2. For analyses, the sequence reads mapping equine genome were removed and the remaining reads were assembled with de novo assembly (MIRA 5 software *(3)*). Because the data contained very little poxvirus sequence (mostly consisting of partial late transcription factor VTFL-1 (ORF045) and envelope phospholipase (ORF011) as well as smaller amounts of partial protein kinase (ORF130), and DNA-directed RNA polymerase subunit RPO132 (ORF101)), no further analysis was done, and we decided to use an alternative approach.

Table A2: Primers used in the first sequencing attempt

| Primer name | Sequence | Target | Ref |
| --- | --- | --- | --- |
| PPP1 | GTCGTCCACGATGAGCAGCT | Major envelope protein of parapoxviruses/ envelope phospholipase (ORF011) | (*2)* |
| PPP4 | TACGTGGGAAGCGCCTCGCT |  |  |
| Pan-pox high GC For | CATCCCCAAGGAGACCAACGAG | RNA polymerase of poxviruses with >60% GC-content (ORF056) | (*4*) |
| Pan-pox high GC Rev | TCCTCGTCGCCGTCGAAGTC |  |  |
| Parapox 045F | CCTACTTCTCGGAGTTCAGC | Late transcription factor 1 of Orf virus (ORF045) | (*5*) |
| Parapox 045R | GCAGCACTTCTCCTCGTAG |  |  |
| ATI up1 | AATACAAGGAGGATCT | A-type inclusion body protein gene of orthopoxviruses (ORF103-104) | (*6*) |
| ATI low1 | CTTAACTTTTTCTTTCTC |  |  |
| Pan-pox low GC For | ACACCAAAAACTCATATAACTTCT | Metalloprotease | (*4*) |
| Pan-pox low GC Rev | CCTATTTTACTCCTTAGTAAATGAT | gene of orthopoxviruses (ORF037) |  |
| A3LFor1 | CNTCHACNMABRAYTGG | P4b precursor gene of chordopoxviruses (ORF079)* | (*7*) |
| A3LRev3 | TGYTCYTCRTCNGHCAT |  |  |
| F10LF958 | GAYYTNAARCCNGAYAA | Protein kinase gene of chordopoxviruses (ORF130)* | (*8*) |
| F10LR1167 | AARTGRAARTCRTARWACCA |  |  |

*Degenerate primers, which have been used to successfully amplify PCR products from distinct squirrel poxviruses

PCR-free sequencing and sequence analysis

To separate host and virus DNA, methylated DNA was removed from 1 µg of DNA with Microbiome DNA Enrichment kit (NEBNext) according to the manufacturer’s instructions. Unmethylated DNA was then subjected to next-generation sequencing (NGS) as described for the first protocol. Before proceeding with the sample, the protocol was tested with 1 µg of cultured reindeer pseudocowpox virus (F00.120R) DNA, from which the entire genome was acquired (assembly performed with Bowtie 2 (*9*) and GQ329669.1 as a reference genome (*10*)).

We were not able to perform a reference-based assembly since no complete genomes of similar viruses are available. Single-end reads were preprocessed and assembled using Lazypipe 2.0 (*11*) with default options and no host filtering. For assembler we used megahit version 1.2.9 (*12*) with the help of in-house scripts.

Sequence alignment, p-distance calculations, and GC-content calculation were performed with MEGA 11 (*13*). Sequences were aligned with Muscle (*14*). A representative of each genus in the subfamily *Chordopoxvirinae* was selected for a phylogenetic analysis of the DNA polymerase gene. Additionally, representatives of each recognized parapoxvirus (PPV) species with complete genome available as well as Molluscum contagiosum virus and Squirrelpox virus as outgroups were selected for phylogenetic analyses of PPVs using the DNA polymerase, early transcription factor, RNA polymerase, and topoisomerase 1 genes. Phylogenetic analyses were performed using amino acid sequences (DNA polymerase) or nucleotide sequences (other genes) using IQ-TREE 2.2.0.3 (*15*) and visualized with iTOL (*16*).

Virus culturing

In an attempt to isolate the virus, the remaining skin biopsy sample (approximately 10 mg) was homogenized in 500 µl of Dulbecco’s phosphate buffered saline + 0.2% bovine serum albumin with a mortar on dry ice. The sample suspension was filtered through 0.45 µm filter (Whatman) to get rid of bacteria. Primary bovine esophagus cells (CCLV, Friedrich-Loeffler-Institute) cultured in Minimum Essential Medium (Gibco) supplemented with 1% L-glutamine (Gibco), 1% Non-essential amino acids (Gibco), 10% sheep serum, 200 IU/ml penicillin (Orion) and 400 µg/ml streptomycin (Sigma) were used in virus culturing. Cells that had been divided on the previous day were infected with 50 µl of the sample supernatant in a 24-well plate (Nunc). The following passages were infected with 1 ml of the previous passage in a 25-cm^2^ cell culture flask (Nunc). Samples were allowed to adsorb for 1 h at 37°C before adding the cell culture medium. Cells were harvested on the seventh day after infection by freezing and thawing three times. The passaging was repeated three times. No cytopathic effect was detected.

Table A3: Amino acid identities (as percentages) of the DNA polymerase gene between F14.1158H, parapoxviruses, and representatives of other Chordopoxviruses. Identities within genus are highlighted with gray.

|  | SPPV | SWPV | CRV | FWPV | YMTV | SQPV | DPV | MYXV | EMCLV | MOCV | CPXV | VACV | MPXV | PCPV/F00.120R | PCPV/VR634 | GSEPV | RDPV | ORF/NZ2 | ORF/HSN-20 | BPSV |
| --- | --- | --- | --- | --- | --- | --- | --- | --- | --- | --- | --- | --- | --- | --- | --- | --- | --- | --- | --- | --- |
| MZ682626/SWPV/Suipoxvirus | 75 |  |  |  |  |  |  |  |  |  |  |  |  |  |  |  |  |  |  |  |
| NC_008030/CRV/Crocodylidpoxvirus | 45 | 45 |  |  |  |  |  |  |  |  |  |  |  |  |  |  |  |  |  |  |
| NC_002188/FWPV/Avipoxvirus | 53 | 54 | 47 |  |  |  |  |  |  |  |  |  |  |  |  |  |  |  |  |  |
| NC_005179YMTV/Yatapoxvirus | 70 | 68 | 45 | 52 |  |  |  |  |  |  |  |  |  |  |  |  |  |  |  |  |
| NC_022563SQPV/Sciuripoxvirus | 58 | 57 | 50 | 50 | 59 |  |  |  |  |  |  |  |  |  |  |  |  |  |  |  |
| NC_006966/DPV/Cervidpoxvirus | 76 | 78 | 45 | 53 | 71 | 60 |  |  |  |  |  |  |  |  |  |  |  |  |  |  |
| NC_001132/MYXV/Leporipoxvirus | 75 | 76 | 46 | 52 | 70 | 59 | 77 |  |  |  |  |  |  |  |  |  |  |  |  |  |
| MN339351/EMCLV/Molluscipoxvirus | 51 | 52 | 53 | 51 | 51 | 59 | 53 | 52 |  |  |  |  |  |  |  |  |  |  |  |  |
| MH320556/MOCV/Molluscipoxvirus | 51 | 52 | 52 | 52 | 50 | 59 | 53 | 51 | 73 |  |  |  |  |  |  |  |  |  |  |  |
| NC_003663/CPXV/Brighton_Red/Orthopoxvirus | 66 | 66 | 47 | 51 | 67 | 61 | 68 | 67 | 54 | 54 |  |  |  |  |  |  |  |  |  |  |
| NC_006998/VACV/West_Reserve/Orthopoxvirus | 66 | 66 | 48 | 51 | 67 | 61 | 68 | 67 | 54 | 53 | 99 |  |  |  |  |  |  |  |  |  |
| ON959143/MPXV/MPX-96/Orthopoxvirus | 66 | 65 | 47 | 51 | 66 | 60 | 67 | 66 | 54 | 53 | 98 | 98 |  |  |  |  |  |  |  |  |
| GQ329669/PCPV/F00.120R/Parapoxvirus | 54 | 54 | 48 | 48 | 53 | 59 | 54 | 55 | 55 | 53 | 57 | 57 | 56 |  |  |  |  |  |  |  |
| GQ329670/PCPV/VR634/Parapoxvirus | 54 | 54 | 49 | 48 | 54 | 60 | 54 | 54 | 55 | 53 | 56 | 57 | 56 | 98 |  |  |  |  |  |  |
| KY382358/GSEPV/Parapoxvirus | 55 | 54 | 48 | 47 | 54 | 58 | 55 | 54 | 54 | 52 | 56 | 56 | 56 | 84 | 84 |  |  |  |  |  |
| KM502564/RDPV/Parapoxvirus | 53 | 52 | 48 | 46 | 52 | 58 | 54 | 53 | 55 | 53 | 56 | 56 | 55 | 85 | 86 | 85 |  |  |  |  |
| DQ184476/ORF/NZ2/Parapoxvirus | 54 | 54 | 49 | 47 | 54 | 59 | 54 | 55 | 55 | 53 | 57 | 57 | 56 | 95 | 94 | 84 | 86 |  |  |  |
| MW537048/ORF/HSN-20/Parapoxvirus | 54 | 54 | 48 | 47 | 54 | 59 | 54 | 55 | 55 | 53 | 57 | 57 | 56 | 95 | 94 | 85 | 86 | 99 |  |  |
| NC_005337/BPSV/Parapoxvirus | 54 | 54 | 48 | 48 | 54 | 59 | 54 | 55 | 54 | 53 | 57 | 57 | 56 | 88 | 88 | 85 | 87 | 87 | 87 |  |
| F14.1158H | 53 | 53 | 50 | 46 | 52 | 60 | 54 | 53 | 56 | 54 | 57 | 57 | 56 | 79 | 80 | 76 | 78 | 79 | 79 | 78 |

CRV = Crocodilepox virus, EMCLV = Equine molluscum contagiousum-like virus, MOCV = Molluscum contagiousum virus, SQPV = Squirrelpox virus, CPXV = Cowpox virus, MPXV = Monkeypox virus, VACV = Vaccinia virus, YMTV = Yaba monkey tumor virus, SPPV = Sheeppox virus, MYXV = Myxoma virus, SWPV = Swinepox virus, DPV = Deerpox virus, FWPV = Fowlpox virus
